# Spatiotemporal bioprinting of microtissues and growth factors within a support bath to engineer anisotropic, zonally defined meniscal grafts

**DOI:** 10.64898/2026.07.22.740001

**Authors:** Francesca D. Spagnuolo, Gabriela S. Kronemberger, Daniel J. Kelly

**Affiliations:** Trinity Centre for Biomedical Engineering, Trinity Biomedical Sciences Institute, Trinity College Dublin, Dublin, Ireland; Department of Mechanical, Manufacturing and Biomedical Engineering, School of Engineering, Trinity College Dublin, Dublin, Ireland; Department of Anatomy and Regenerative Medicine, Royal College of Surgeons in Ireland, Dublin, Ireland; Advanced Materials and Bioengineering Research Centre (AMBER), Royal College of Surgeons in Ireland and Trinity College Dublin, Dublin, Ireland

**Keywords:** microtissues, bioprinting, support bath, growth factors, meniscus

## Abstract

Current clinical treatments for meniscal injuries remain limited and are associated with an increased risk of developing osteoarthritis (OA). This has motivated the development of tissue engineering (TE) strategies to engineer more biomimetic meniscal grafts capable of promoting functional joint regeneration. Existing approaches typically fail to recapitulate the zonal heterogeneity of the native meniscus, which contains distinct inner and outer regions with unique extracellular matrix (ECM) composition and organization. Here, we introduce a novel bioprinting strategy using spatially patterned growth factors and mesenchymal stromal/stem cell (MSC)-derived microtissues (µTs) to engineer meniscal constructs with zonally defined structure and composition. We first investigated the effects of different growth factor regimes, specifically connective tissue growth factor (CTGF) and transforming growth factor-β3 (TGF-β3), on fibrochondrogenesis of MSC-derived µTs. While TGF-β3 alone promoted a more inner-zone meniscus phenotype, stimulation of µTs with a combination of TGF-β3 and CTGF supported the development of tissues that more closely mimicked the outer zone of the meniscus. Using laponite to control the release of these growth factors, it was also possible to bioprint zonally defined meniscal tissue within a methacrylate xanthan gum (XG-MA) support bath. A ‘fibro-ink’ containing µTs, CTGF and TGF-β3 supported higher collagen type I deposition and lower collagen type II deposition, while a ‘chondro-ink’ containing µTs and TGF-β3 promoted higher collagen type II deposition. Based on these findings, dual-cartridge bioprinting was next used to spatially pattern µTs with CTGF + TGF-β3 (fibro-ink) or TGF-β3 (chondro-ink) to generate regionally defined, meniscal-like engineered tissues. This approach enabled the bioprinting of scaffold-free constructs with aligned collagen and zone-specific ECM depositions, with an inner region consisting of sGAG and collagen types I and II, and an outer region rich in sGAG and collagen type I. These findings highlight the potential of co-printing both growth factors and MSC-derived µTs for engineering scaffold-free, zonally defined meniscal tissues.

## 1. Introduction

Meniscal injuries are a major cause of degenerative changes in the knee joint and are strongly associated with the onset of OA [1–3]. Since the meniscus plays a critical role in load distribution and joint stability, its impairment can lead to significant functional limitations, even during low-impact activities such as walking [4, 5]. However, the regenerative capacity of the meniscus is very limited, which greatly increases the risk of OA following injury [6]. Current clinical treatments, including partial meniscectomy and meniscal allograft transplantation (MAT), remain the gold standards for managing meniscal injuries [7–9]. Nevertheless, their long-term outcomes are often unsatisfactory, as these strategies fail to fully restore the native meniscal structure and function. This limitation has driven growing interest in musculoskeletal TE approaches to develop biomimetic meniscal replacements.

The meniscus is a crescent-shaped fibrocartilaginous tissue composed of two distinct zones: an inner avascular region, rich in sulfated glycosaminoglycans (sGAGs) and type II collagen, and an outer vascularized region, characterized by a lower sGAG content and a higher amount of type I collagen[10, 11]. Overall, the tissue contains elevated levels of type I collagen, which is highly organized, consisting of both circumferentially and radially aligned fibers, conferring strong anisotropy to the meniscus[12]. Early attempts at engineering functional meniscal grafts commonly relied on meniscus-derived cells, enzymatically isolated from the native tissue and either seeded into scaffolds or hydrogels, or allowed to self-assemble to generate meniscus-like constructs [13–15].

While these studies were highly informative, the resulting tissues typically failed to mimic the zonal composition and anisotropic architecture of the native meniscus. In an attempt to address this challenge, poly (lactic-co-glycolic acid) (PLGA) microparticles have been used for region-specific delivery of growth factors (CTGF and TGF-β3) to promote the development of distinct meniscal zones [16–18]. In preclinical models, scaffolds enriched with such PLGA microparticles promoted meniscus regeneration with zonally distinct features, specifically an inner region rich in sGAGs and collagen type II, and an outer region rich in collagen type I [18]. This regenerative approach required the recruitment of endogenous progenitor cells from the joint space into the scaffold to promote meniscal tissue formation. In another study, the local delivery of TGF-β3 and basic fibroblast growth factor (bFGF) was explored to support the integration and maturation of a polycaprolactone (PCL) scaffold for meniscus repair, where TGF-β3 controlled matrix production and bFGF supported cellular recruitment to the defect site[19]. While conceptually appealing, it remains unclear whether such cell-free approaches will be as efficacious in the human clinical population, where endogenous cells may be incapable of mounting a similar regenerative response. Regardless, these previous studies point to the clear benefits of controlled growth factor presentation for meniscus regeneration, which could potentially be integrated into an exogenous cell-based, meniscus TE strategy to generate more regionally defined grafts.

Bottom-up TE is a relatively new paradigm that relies on assembling cellular building blocks, such as cell sheets, spheroids, or µTs, to generate scaled-up constructs. µTs are particularly advantageous because they act as single, functional tissue units that can be assembled without the need for exogenous scaffolds, thereby avoiding complications associated with implantation of biomaterials into the body[20, 21]. We have previously demonstrated that region-specific fibrocartilage grafts can be engineered by bioassembling two distinct populations of µTs generated using meniscus progenitor cells (MPCs): one population derived from inner zone MPCs and the other from outer zone MPCs. The resulting graft consisting of an inner zone rich in sGAGs and collagen type I and II, and an outer zone consisting of sGAGs and primarily collagen type I [22]. However, isolating and expanding distinct MPC populations to engineer patient-specific autografts remains technically and logistically challenging, impeding the translation of such approaches into a clinical setting. Mesenchymal stem/stromal cells (MSCs) offer a more clinically feasible alternative cell source due to their easier accessibility and multipotency [23, 24]. Furthermore, we have recently shown that MSC derived µTs can generate structurally aligned meniscus-like tissue when bioprinted into support baths with a stiffness supportive of a fibrochondrogenic phenotype [25]. If combined with a strategy to promote zone-specific meniscus phenotypes, such a bioprinting platform could potentially be used to engineer anisotropic, zonally defined meniscal grafts.

The goal of this study was to bioprint a gelatin-laponite-based bioink containing MSC derived µTs into a mechanically defined methacrylate Xanthan Gum (XG-MA) support bath to engineer regionally defined meniscal grafts. Laponite was introduced into the bioink to support the spatiotemporally defined release of CTGF and TGF-β3, with the goal of engineering zonally defined meniscal grafts. Nanosilicates such as laponite have been widely applied in TE and 3D bioprinting [26, 27]. Owing to their positive surface charge, laponite nanosilicates can readily bind growth factors and have previously been leveraged to develop composite inks for 3D bioprinting with controlled protein release profiles [28]. In the first phase of this study, µTs were subjected to different growth factor stimulation regimes, specifically CTGF and/or TGF-β3 stimulation. The growth factor combinations supportive of either an inner or outer zone meniscus phenotype were then integrated into the gelatin-laponite-µT inks and used to bioprinted filaments of aligned meniscal tissues. Finally, we sought to use dual-cartridge bioprinting, specifically with one bioink loaded with µTs and TGF-β3 for the inner zone (termed ‘chondro-ink’), and another with µTs, CTGF and TGF-β3 for the outer zone (termed ‘fibro-ink’), to fabricate scaled-up, region-specific fibrocartilage grafts within a XG-MA support bath. In doing so we sought to address the multiple challenges of engineering scaled-up meniscal tissues with zonally defined structure and composition.

## 2. Materials and Methods

### 2.1 Isolation and expansion of gBM-MSCs and µT formation

Bone marrow-derived MSCs were isolated from the sternum of a skeletally mature female goat. The extracted marrow was washed in expansion medium containing high-glucose Dulbecco’s Modified Eagle Medium (hgDMEM), 10% fetal bovine serum (FBS) and penicillin (100 U/mL) – streptomycin (100 μg/ml) (all from Gibco) and triturated with a 16G needle until a homogenous mixture was obtained. The suspension was then centrifuged at 650 g for 5 min, and the resultant cell pellet was resuspended in fresh expansion medium twice before it was filtered through a 40 μm cell sieve (Sarstedt). Cell counting was performed with trypan blue in the presence of acetic acid (6% final) before plating at a density of 1.3 × 10^5^ cells/cm^2^. Following colony formation, cells were trypsinized, counted, and plated for 2 additional passages at a density of 5 × 103 cells/cm^2^ at 5% pO2 in expansion medium supplemented with 5 ng/ml of fibroblast growth factor (FGF)-2 (PeproTech Ltd). Medium change was performed three times per week. The µTs were fabricated as previously described (Chapter III). The 3D printed mold stamps containing 1,889 micro resections and 4% (w/v) ultrapure agarose were sterilized in an autoclave at 120°C for 20 minutes. Next, 6 mL of ultrapure agarose was gently poured into each well of the 6-well plate and allowed to cool down for 15 min after molding it. Agarose molds were then soaked overnight in XPAN, at 37°C in a humidified atmosphere with 5 % CO2 and 5 % O2. MSCs were seeded at a density of 7,55×10^5^ into the microwells in the final volume of cell suspension of 5 mL of CDM+. 20 minutes after seeding, the plates were centrifuged for 5 minutes at 700 x g to collect cells at the bottom of each well. After 2 days, µTs were harvested as described before[25] [29], and used at a density of 45,000 µTs/mL for bioprinting experiments.

### 2.2 µT fusion and fibrochondrogenic differentiation of fused µTs

On day 2, 12-well plates were coated with 2% (w/v) sterile, autoclaved agarose and equilibrated with XPAN for 1 h. For each condition, 150 µTs were seeded per well and incubated at 37 °C for 1 h to pre-fuse before gently adding medium. Cultures were maintained for 4 weeks in CDM− basal medium with the following regimens: (1) TGF-β3, 10 ng/mL (4 weeks); (2) CTGF, 100 ng/mL (4 weeks); (3) TGF-β3 → CTGF, 10 ng/mL TGF-β3 for 2 weeks followed by 100 ng/mL CTGF for 2 weeks; (4) CTGF → TGF-β3, 100 ng/mL CTGF for 2 weeks followed by 10 ng/mL TGF-β3 for 2 weeks; (5) CTGF + TGF-β3, 100 ng/mL CTGF plus 10 ng/mL TGF-β3 for 4 weeks. Medium was changed three times per week.

### 2.3 Support bath (XG-MA) and bioink synthesis (laponite/gelatin)

XG (0.5% w/v) in ultrapure water reacted with glycidyl methacrylate (40 mL) overnight at 60 °C in the dark, dialyzed 7 days (twice-daily water changes), lyophilized 48 h, stored at −20 °C, and EO-sterilized (12 h). Two days before printing, XG-MA was dissolved in phenol red-free DMEM with penicillin (100 U/mL) and streptomycin (100 µg/mL) and rotated at 40 rpm at room temperature. On the print day, LAP was added (0.25% w/v), the bath was centrifuged to de-gas (2,500 × g, 5 min, RT), and 4 mL was dispensed into a 6-well plate using a positive-displacement pipette.

For the bioink preparation, Laponite XLG (BYK Additives & Instruments, UK) was autoclaved at 120 °C for 20 min and subsequently dried to remove residual moisture. DMEM was then added, and the suspension was stirred at 700 rpm for 2 days on a magnetic stir plate. The laponite dispersion was combined with a sterile gelatin solution and the appropriate growth factor, then stirred overnight at room temperature to promote growth factor binding to the nano clay and to obtain a homogeneous mixture. On the day of printing, the bioink was chilled at 4 °C to promote complete gelation and then mixed with µTs to yield final concentrations of 1% (w/v) gelatin and 0.05% (w/v) laponite and 45,000 µTs/mL. The total growth factor load was calculated as the cumulative dose for a 4-week study, assuming 250 mL of medium: 25 µg for CTGF and 2.5 µg for TGF-β3.

### 2.4 Rheological analysis of bioinks

Rheological tests were performed on a MCR 102 rheometer (Anton Paar) with Peltier temperature control using a 25 mm parallel-plate geometry (PP25). The flow curve (viscosity vs shear rate, 0.1-1000 s⁻¹ was acquired at 4°C for laponite/gelatin-based bioinks. Samples were maintained in a high-humidity environment to prevent dehydration. All measurements were performed in triplicate.

### 2.5 Growth factor release from bioprinted filaments in media and release quantification over time

To evaluate 4-week release profiles of CTGF and TGF-β3 bound to laponite, bioinks loaded with either factor were printed into a 1% XG-MA support bath and UV-crosslinked for 4 min (as described previously). For each well of a 6-well plate, eight 10 mm filaments were printed, corresponding to a theoretical load of 0.8 µg TGF-β3 per well or 8 µg CTGF per well. Each condition was printed in triplicate. Constructs were cultured for 28 days; medium was sampled twice per week and snap-frozen at −70 °C. Growth factor concentrations in the collected media were quantified by ELISA for CTGF and TGF-β3 (Bio-Techne), following the manufacturer’s protocol, with absorbance read at 450 nm on a microplate reader. For analysis, samples collected within the same nominal interval were pooled and assigned to the terminal day of that interval (e.g., days 2, 5, and 7 pooled and reported as day 7).

### 2.6 Bioprinting of µTs and dual cartridge bioprinting set-up

For bioprinting, µTs were harvested on day 2 and mixed with the designated bioinks. For single-filament prints, µTs were combined with three formulations: (1) TGF-β3-loaded bioink (2.5 µg), (2) dual-loaded bioink containing CTGF and TGF-β3 (25 µg and 2.5 µg, respectively), and (3) a non-loaded bioink, with the growth factor supplied in the culture medium changed three times per week. For prints in which the growth factor was incorporated into the ink, the culture medium was basal CDM−. For dual-cartridge prints. Two cartridges were used: a “chondro-ink” (TGF-β3, 2.5 µg) and a “fibro-ink” (CTGF plus TGF-β3). Both syringes were prepared, loaded into the bioprinter (CELLINK), and calibrated to the same origin. For the meniscus construct, three half-circumferential lines were printed for the outer region and two lines for the inner region. Printing employed a piston-based syringe-pump printhead with an extrusion rate of 6 µL/s and a print speed of 2 mm/s.

### 2.7 Histological analysis

For construct analysis, constructs were retrieved by mechanically disrupting the surrounding support bath and carefully extracting the samples using tweezers Briefly, samples were fixed in 4% paraformaldehyde (PFA) overnight at 4 °C. After fixation, they were dehydrated through a graded ethanol series (50–100% v/v), cleared in xylene, and embedded in paraffin wax (Sigma) as previously described. Sections of 5 µm thickness were cut, rehydrated, and subjected to histological staining. Haematoxylin and eosin (H&E; Sigma) were used to assess tissue morphology, ECM, and nuclei. sGAG content was stained with 1% (w/v) Alcian Blue (AB) 8GX in 0.1 M HCl and counterstained with 0.1% (w/v) nuclear fast red for cell distribution. Collagen deposition was visualized using 0.1% (w/v) Picrosirius Red (PR; Sigma). Stained sections were mounted in Pertex (Avantor) and imaged with an Aperio ScanScope slide scanner (Leica, Germany).

### 2.8 Biochemical evaluation

After retrieval, the samples were washed twice with 1X PBS and dried. Papain enzyme solution created with 3.88 U/mL of papain enzyme, 100 mM sodium phosphate buffer, 5 mM Na2EDTA, and 10 mM L-cysteine at pH 6.5 (all from Sigma), was then added to digest the samples at 60°C for 18 hours. Immediately after digestion, DNA content was quantified using the Quant-iT™ PicoGreen® dsDNA Reagent and Kit (Molecular Probes, Biosciences). The amount of sulphated glycosaminoglycan (sGAG) was determined using a dimethylmethylene blue dye (DMMB, Sigma) binding assay. To exclude any background absorbance, the pH of the DMMB was adjusted to 1.5. A chondroitin sulphate standard (1 mg/mL, Sigma) was used, and the absorbance was measured at 530 and 590 nm using a Synergy HT multi-detection microplate reader (BioTek Instruments, Inc). The 530/590 absorbance ratio was used to generate the standard curve and determine the sGAG concentration from the digested sample. Total collagen content was determined by measuring the hydroxyproline content using a chloramine-T assay. The samples were mixed with 38% HCl (Sigma) (1:1 ratio) and incubated at 110°C for 18 hours for hydrolysis. After cooling, samples were centrifuged at 5000 x g for 5 minutes, and let dry at 60°C for 48 h. The sediment was reconstituted in 200 µL of ultra-pure H2O. Chloramine T (2.82% w/v) and 4-(Dimethylamino) benzaldehyde (0.05% w/v) were added and incubated for 20 minutes at RT in the dark. The hydroxyproline content was quantified using a trans-4-Hydroxy-L-proline standard at a wavelength of 570 nm with a Synergy HT multi-detection microplate reader (BioTek Instruments, Inc). Hydroxyproline levels were estimated from the standard curve at a wavelength of 570 nm. The hydroxyproline-to-collagen ratio of 1:7.69 was used to calculate the collagen content.

### 2.9 RT-qPCR analysis of bioprinted constructs

At day 28 post-bioprinting, the samples were washed 3 times with 1X PBS and then snap frozen for further processing. RNA was isolated from the samples using the Trizol method (Sigma-Aldrich), following the manufacturer’s instructions. Chloroform extraction was performed, and the RNA was resuspended in RNase-free water. The RNA samples were then stored at -80°C. For cDNA synthesis, a High-Capacity RNA-to-cDNA™ Kit (Applied Biosystems™), was used to transcribe 500 ng of RNA from each sample into cDNA (20 µL) via polymerase chain reaction (PCR). Real-time PCR was performed using an Applied Biosystems instrument, and Taqman PCR universal master mix (ThermoFisher) was used for the PCR reaction. Human-specific TaqMan probes (ThermoFisher) were used for gene amplification. The qPCR amplification followed the following cycles: 50°C for 2 minutes, 95°C for 10 minutes and 40 cycles at 60°C for 1 minute each. The levels of gene expression were quantified using the real-time PCR data and analyzed with the 2(-ΔΔCt) method. The expression levels were normalized to the average expression of the 18S rRNA housekeeping gene. All experiments were performed in triplicate.

### 2.10 Live/Dead staining

Cell viability was assessed using a live/dead assay kit (Invitrogen, Bioscience). The constructs were first rinsed in PBS 1X and then incubated for 1 hour in a solution containing 2 μM of calcein and 4 μM ethidium homodimer-1 (EthD-1). Following the incubation period, the constructs were rinsed again to remove any excess of the dyes and were imaged using a Leica SP8 scanning confocal microscope. Excitation wavelengths of 485 nm and 530 nm were used for calcein, while emission wavelengths of 530 nm and 645 nm were used for calcein and EthD-1. Maximum projection z-stack reconstructions were generated to analyze cell viability throughout the depth of the tissue, with the view from the top.

### 2.11 Immunofluorescence evaluation

Following fixation in 4% paraformaldehyde overnight and paraffin processing (Section 6.2.5), sections were rehydrated through graded ethanol (100–50%) into 1× PBS (Gibco). Antigen retrieval was performed enzymatically with Hyaluronidase (4,000 U/mL, 25 min, 37 °C; Sigma-Aldrich) followed by Pronase (3.5 U/mL, 25 min, 37 °C, Sigma). Sections were rinsed and blocked for 1 h at room temperature in PBS containing 1% BSA and 10% goat serum. Primary antibodies to collagen I (Abcam, ab90395; 1:400) and collagen II (Santa Cruz, sc-52658; 1:400) were applied in blocking buffer and incubated overnight at 4 °C in a humidified chamber. After three PBS washes, appropriate secondary antibodies in 2% BSA were applied for 1 h at room temperature in the dark. Slides were washed, mounted with Fluoroshield containing DAPI (Sigma), sealed, and stored at 4 °C in the dark. Imaging was performed on a Leica SP8 confocal microscope; representative images are shown as maximum-intensity projections of z-stacks.

### 2.12 Image quantification analysis of immunofluorescence pictures and PLM analysis

For immunofluorescence quantification, the intensity of 3 specific regions of interest (ROI) within each sample was measured and quantified through ImageJ software. The intensity was then normalized by the number of nuclei in the ROI. For PLM analysis, Sections were stained with 0.1% (w/v) Picrosirius Red, mounted with DPX (Sigma-Aldrich), and imaged on an Olympus BX41 polarizing light microscope equipped with a MicroPublisher 6 CCD camera and an Olympus U-CMAD3 adapter. Collagen fibril orientation and coherency were quantified in ImageJ using the OrientationJ and Directionality plugins.

## 3. Results

### 3.1 MSC derived µTs display distinct patterns of growth and differentiation following stimulation with TGF-β3 and/or CTGF

We first sought to explore how different growth factor stimulation regimes would influence the fusion and (fibro)chondrogenic differentiation of MSC derived µTs (Fig. 1A). TGF-β3 is known to promote chondrogenesis of MSCs and the development of a tissue rich in sGAGs and type II collagen [30, 31] CTGF promotes type I collagen production, while exposure to CTGF followed by TGF-β3 has been shown to support a more fibrocartilage phenotype characterized by the presence of both collagen type I and II[16, 17]. To assess how these growth factors would influence the phenotype of MSC derived µTs, we first stimulated relatively small numbers of µTs (150 µTs per construct; 2,000 cells/µT; 3 × 10^5^ cells/construct) to five different growth factor regimens (Regime A, B, C, D and E; see Fig. 1A). Microtissues cultured in the microwells for 2 days were harvested and seeded into an agarose mold at uniform density (Fig. 1 A–G). For regime A and B, these µTs were exposed to the same growth factor (TGF-β3 or CTGF) for the remainder of the 28-day culture period. For regime C and D the growth factor was switched after 14 days of culture. For regime E, constructs were continuously exposed to both TGF-β3 and CTGF. For all conditions, fusion was evident within 24 h, and over 21 days, the µTs formed compact tissues (Fig. 1 B).

**Figure 1.**
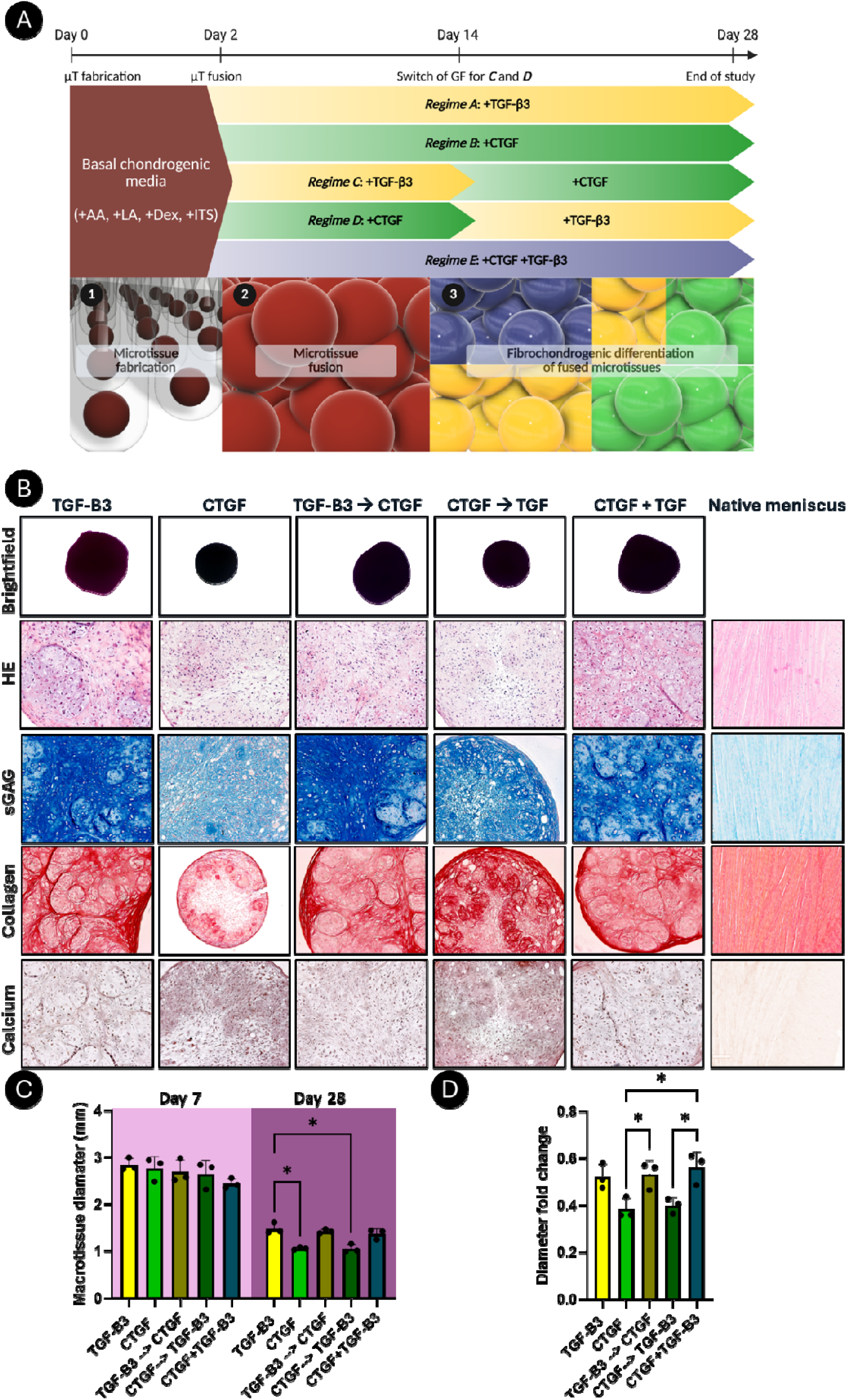
MSC µT fusion and fibrocartilaginous tissue development under altered growth factor stimulation regimes. (A) Schematic of the growth factor regimens applied during µT fusion. µTs were fused at day 2 post formation, and the different culture regimes were applied. At day 14, the growth factors were switched in regimes C and D. At day 28, the study was stopped and the samples analyzed. Step 1: µT fabrication; step 2: µT fusion; step 3: differentiation of fused µTs. (B) Brightfield and histological assessment of fused µTs after 28 days under the following regimens: TGF-β3 for 4 weeks (regime A); CTGF for 4 weeks (regime B); TGF-β3 for 2 weeks followed by CTGF for 2 weeks (regime C); CTGF for 2 weeks followed by TGF-β3 for 2 weeks (regime D); and combined TGF-β3 + CTGF for 4 weeks (regime E). Histological stains: H&E (morphology), Alcian Blue (sGAG), Picrosirius Red (total collagen), and Alizarin Red (calcium deposit/mineralization (C) µT diameter at day 7 and day 28. (D) Diameter fold change across regimens, defined as D28/D1. N=3, significant differences were determined using a two ANOVA followed by a Tukey’s test post hoc comparison for (C) and an ordinary one-way ANOVA followed by a Tukey’s multiple comparison test for (D).

By day 28, construct size differed by regimen, where constructs treated with CTGF for the full 4 weeks and those primed with CTGF followed by TGF-β3 for the subsequent 2 weeks (Fig. 1C, D) were smaller in diameter compared to other groups (p<0.05). Histology demonstrated regimen-dependent sGAG and collagen deposition (Fig. 1B). More intense staining for sGAG deposition was observed in constructs continuously exposed to TGF-β3, as well as in the TGF-β3 to CTGF group. Relatively weak staining for sGAG and collagen deposition was observed in constructs continuously exposed to CTGF (Fig. 1B). Alizarin red staining for calcium deposition was weak in all groups, indicating no mineralization of the resulting tissues (Fig. 1B).

Neo-tissue growth and maturation were evaluated after 4 weeks of culture using a range of biochemical assays (Fig. 2). The DNA content of the CTGF only group was significantly lower than the CTGF+TGF-β3 group (p < 0.05) (Fig. 2 A). After 28 days, all groups containing TGF-β3 contained higher levels of sGAG compared to the CTGF only group, with the greatest accumulation observed in TGF-β3 and CTGF+TGF-β3 groups (Fig. 2 B). This response was maintained when sGAG levels were normalized to DNA (Fig. 2 D). A similar pattern was observed for collagen, with increased deposition observed in groups where TGF-β3 was present during the first two weeks of induction. There was a trend towards higher collagen accumulation in the CTGF+TGF-β3 group compared to TGF-β3 alone (p = 0.07) (Fig. 2 E). However, the CTGF and CTGF→TGF-β3 groups contained significantly less collagen, underscoring the importance of TGF-β3 during the initial induction phase for collagen biosynthesis (Fig. 2 C, E). Normalization of collagen content to DNA yielded similar trends, with statistically significant differences maintained across groups (Fig. 2 E). After 28 days, the wet weight of each construct was also measured (Fig. 2 F). The wet weight of constructs was comparable between TGF-β3 and CTGF+TGF-β3 groups, while it was significantly reduced in the CTGF only group compared to both the TGF-β3 (p<0.05) and CTGF+TGF-β3 groups (p <0.01) (Fig. 2 F). When normalized to wet weight, TGF-β3 stimulation yielded the highest sGAG content (Fig. 2 G), although this value exceeded that typically observed in native meniscus and was more comparable to articular cartilage (Fig. 2 G). In contrast, normalization of collagen to wet weight revealed that the TGF-β3→CTGF and CTGF+TGF-β3 groups contained significantly higher collagen levels than the CTGF and CTGF→TGF-β3 groups (p < 0.01) (Fig. 2 H). Nevertheless, the maximum collagen content achieved (∼8% of wet weight) remained below native meniscus values (∼22%)[32]. These findings highlight the need for a trade-off between maintaining physiologically relevant levels of sGAG and increasing collagen deposition to approach the native tissue composition. Thus, tuning the growth factor stimulation regime allows modulation of matrix composition toward zonal biomimicry of the native meniscus.

**Figure 2.**
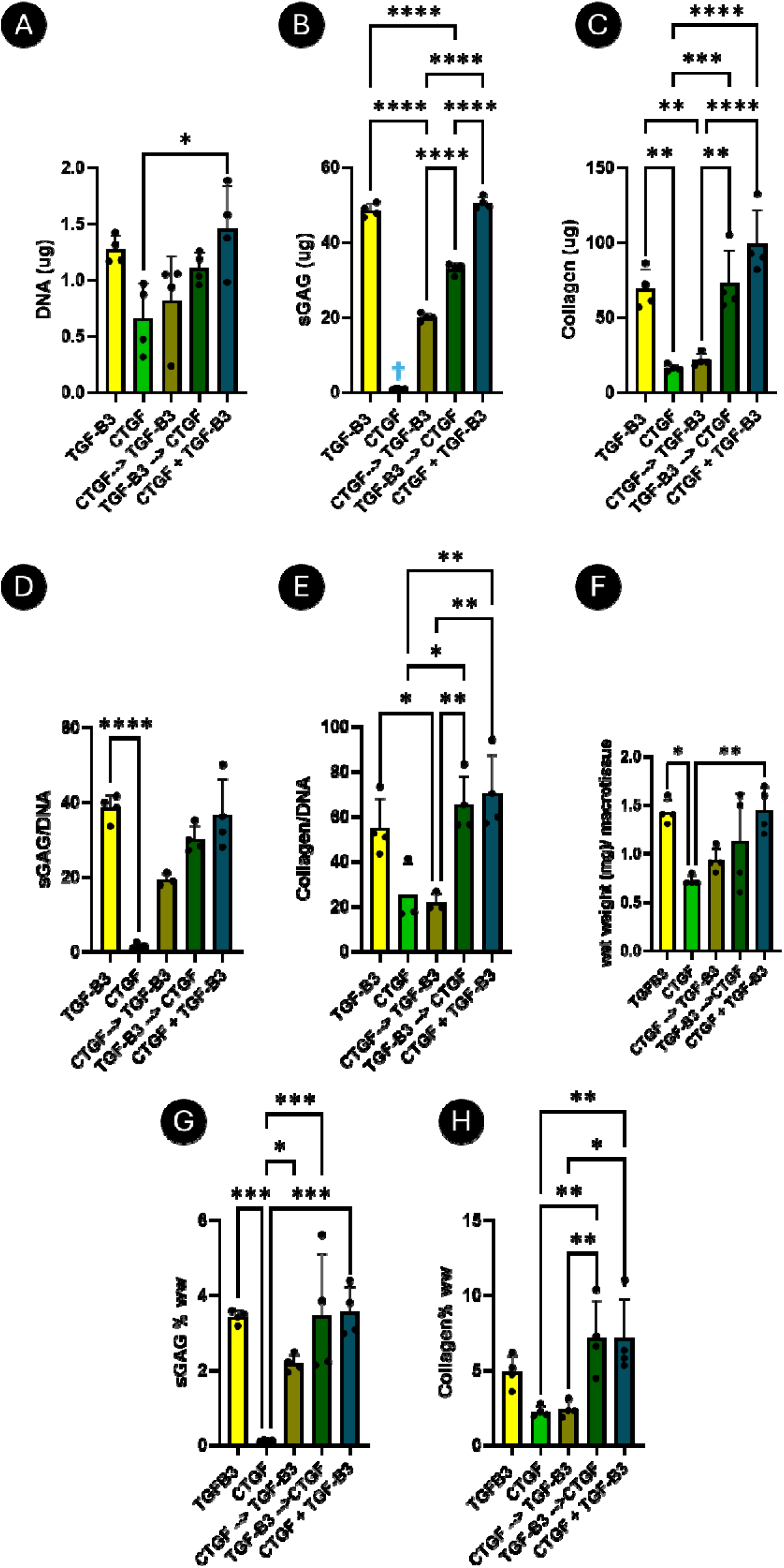
Self-assembled µTs display distinct patterns of growth and matrix deposition after 28 days in culture under different growth factor regimes. (A) DNA quantification, (B) sGAG, (C) Total collagen, (D) sGAG/DNA and collagen/DNA ratios, (F) Wet weight (mg)/µT to compare construct size (G, H) Normalization of sGAG and collagen to wet weight in percentage (%). N=4, significant differences were determined using an ordinary one-way ANOVA followed by a Tukey’s multiple comparison test, where * denotes p < 0.05, ** denotes p < 0.01, *** denotes p < 0.001, and **** denotes p < 0.0001.

After 28 days of culture under different growth factor regimes, fused µTs were analyzed by immunofluorescence to further assess tissue phenotype (Fig. 3). The CTGF+TGF-β3 group stained more intensely for collagen type I deposition (Fig. 3). The TGF-β3 only constructs stained more intensely for collagen type II, consistent with a more hyaline cartilage–like phenotype (Fig. 3). In contrast, CTGF alone, CTGF→TGF-β3, and TGF-β3→CTGF groups exhibited weaker staining for both collagen types (Fig. 3). These findings suggest that combined CTGF+TGF-β3 stimulation best supports an outer zone meniscus phenotype, with collagen type I predominating over collagen type II. Because collagen type II is enriched in the native inner meniscus, two groups were selected for subsequent bioprinting studies: TGF-β3 alone to support an inner region phenotype rich in both collagen type I and II, and CTGF+TGF-β3 to support an outer region phenotype with higher levels of collagen type I (Fig. 3).

**Figure 3.**
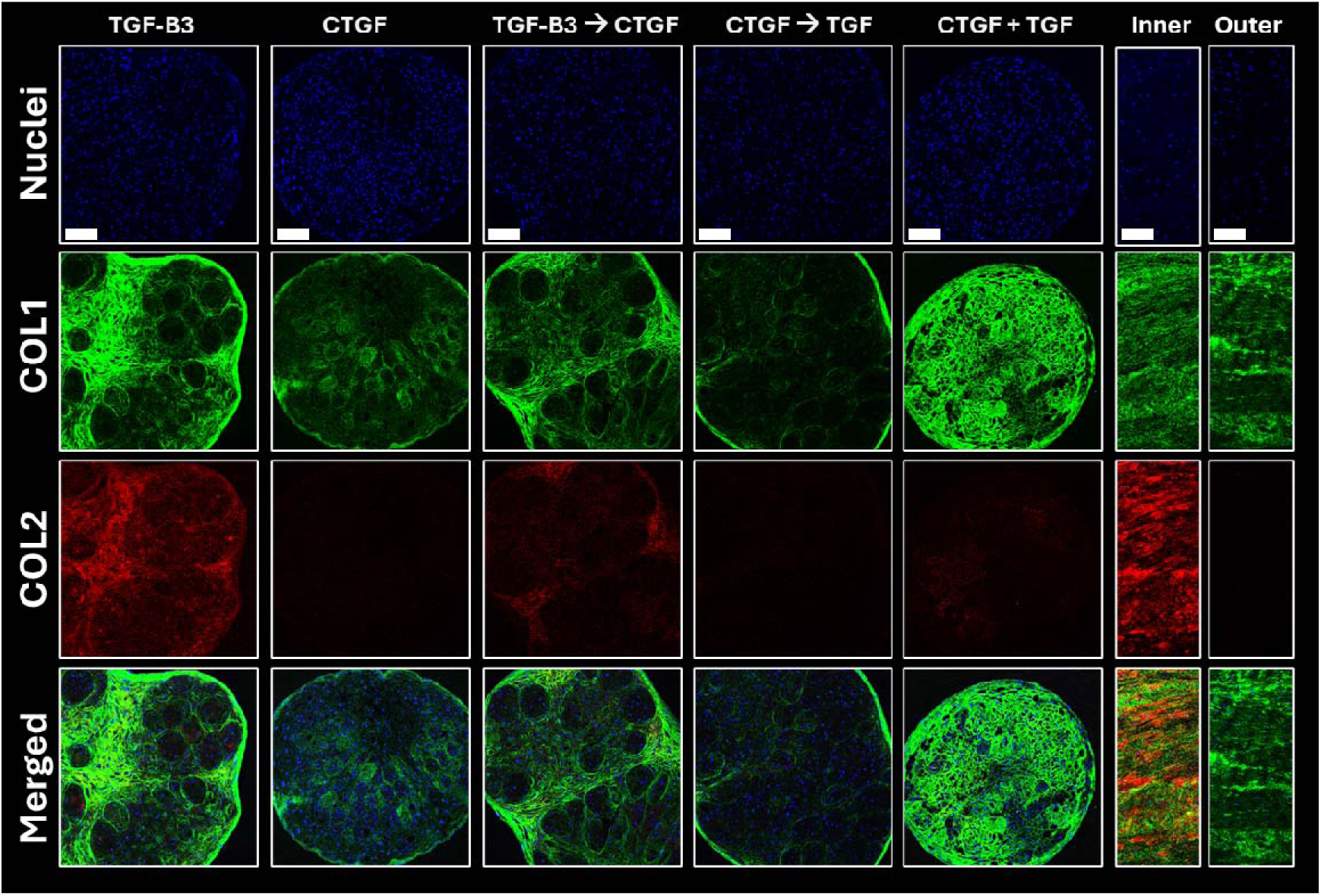
µTs develop a distinct pattern of ECM deposition when exposed to different growth factor regimes. µTs were harvested at 2 days post-fabrication and fused in a low-adherent agarose mold and kept in culture for 28 days at different growth factor regimes: 1) TGF-β3 only for 4 weeks (TGF-β3), 2) CTGF only for 4 weeks (CTGF), 3) TGF-β3 for 2 weeks and switch to CTGF for the remaining two weeks (TGF-β3 ➔ CTGF), 4) CTGF for 2 weeks and subsequently TGF-β3 for the remaining 2 weeks (CTGF➔TGF-β3), 5) CTGF and TGF-β3 together for 4 weeks (CTGF+TGF-β3). At the end of the culture, the resultant grafts were stained against chondrogenic markers, collagen type I (COL1) and collagen type II (COL2). Scale bar: 200µm.

### 3.2 Localized and sustained delivery of growth factors in bioprinted µTs affects tissue compaction and cell viability

After identifying the most suitable growth factor regimens for promoting the zone-specific differentiation of meniscal µT-based constructs, we next sought to determine if such growth factors could be co-printed with µTs within a XG-MA support bath (Fig. 4). The bioink formulation was based on the findings of our previous work, as was the choice of the support bath [25]. Because gelatin functions as a sacrificial bioink, loading growth factors directly into it would result in their rapid release within 24 hours, coinciding with gelatin dissolution [33, 34]. To enable controlled release over four weeks, an additional delivery system was required. Building on previous studies, laponite was incorporated into a gelatin bioink to regulate the release of CTGF and TGF-β3, leveraging the electrostatic interactions between positively charged laponite and the predominantly negatively charged growth factors [35, 36]. This formulation—comprising laponite (0.05 v/w %), gelatin (1%), and µTs (45,000 µTs/mL) —was designated as the principal bioink for the study (Fig. 4 A)

**Figure 4.**
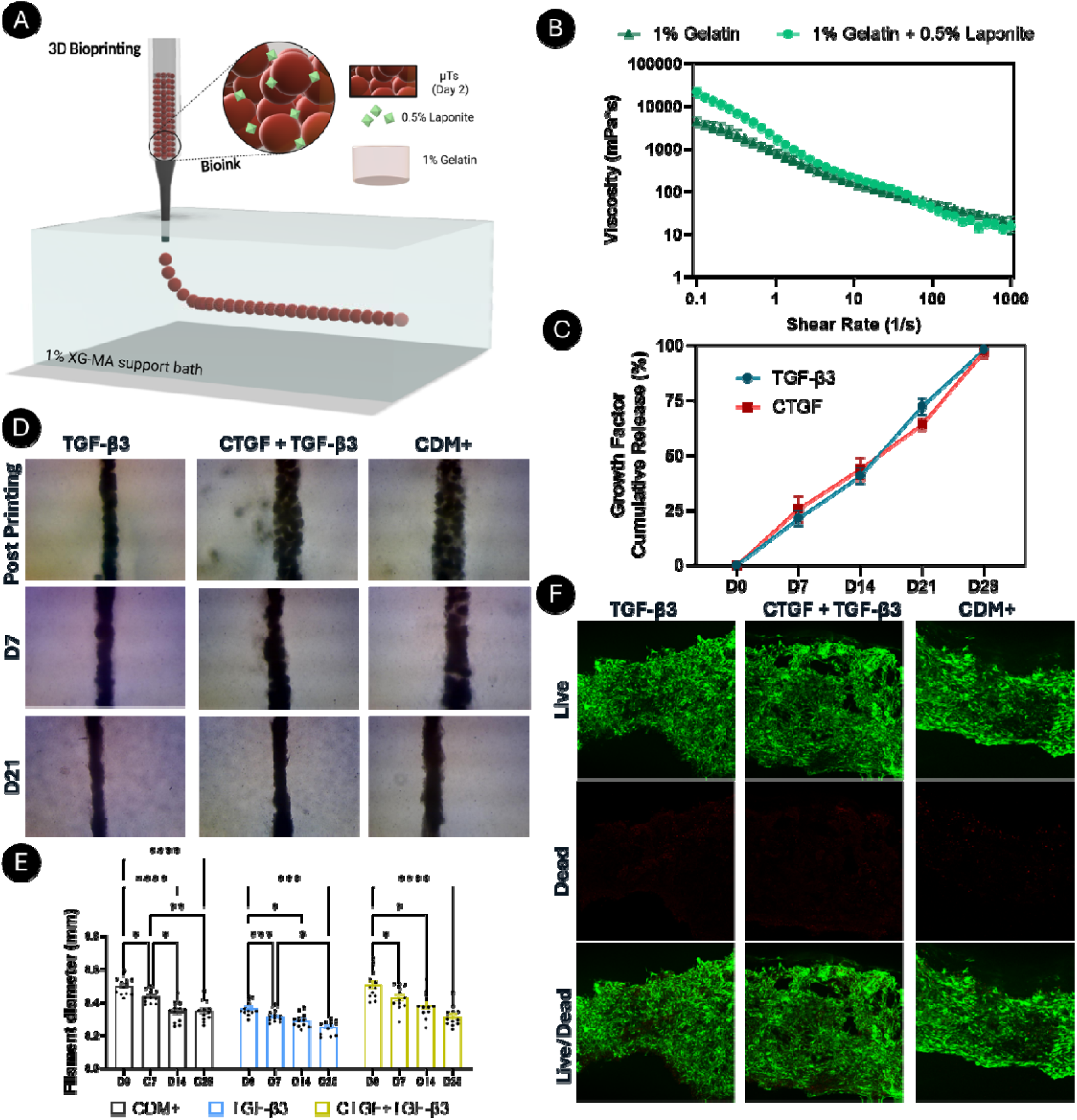
Controlled release of growth factors from laponite functionalized bioinks. A) Schematic representation of the bioink formulation and printing of µTs at a density of 45,000 µTs/mL into an XG-MA support bath. B) Viscosity profiles of gelatin bioink and laponite+ gelatin-based bioink measured over a shear rate range of 0–1000 s⁻¹ at 4 °C (N = 3). C) Release of growth factors from the bioinks assessed by ELISA (N = 3). D) Brightfield microscopy images of printed µTs in fibro-ink supplemented with different growth factors over 21 days of culture, showing tissue compaction and µT fusion. E) Filament diameter of printed µTs in fibro-ink across different conditions (N = 12). F) Viability assessment of printed µTs in fibro-ink after 28 days in culture, post-printing. Significant differences were determined using one-way ANOVA followed by Tukey’s multiple comparison test, where *p < 0.05, **p < 0.01, ***p < 0.001, and ****p < 0.0001.

The influence of incorporating laponite into gelatin on the potential printability of the resulting ink was first evaluated by rheological assessment of their shear-thinning behavior (Fig. 4 B). Both formulations displayed favorable shear-thinning properties, with the laponite-containing bioink exhibiting a higher initial viscosity (Fig. 4B). To characterize the growth factor release kinetics, we quantified the cumulative elution of CTGF and TGF-β3 from the bioink formulations by ELISA at defined time points (Fig. 4 C). Medium collected 6 h after bioprinting (designated day 0) contained no detectable growth factors, confirming the absence of an initial burst release (Fig. 4 C). Thereafter, both factors were released in a sustained manner: cumulative release reached ∼20% by day 7, increased to ∼40–45% by day 14, and to ∼60–70% by day 21. By day 28, cumulative release was 97–100%, indicating near-complete release over the four-week period (Fig. 4 C). The profiles for CTGF and TGF-β3 were comparable throughout, consistent with controlled, gradual delivery from the bioinks.

µTs were then bioprinted into the XG-MA support bath using previously optimized parameters (6 µL/s extrusion rate, 3 mm/s print speed) (Fig. 4 D). Three main groups were investigated: (1) a laponite/gelatin bioink supplemented with TGF-β3, hereafter referred to as the *chondro-ink*; (2) a laponite/gelatin bioink supplemented with CTGF+TGF-β3, referred to as the *fibro-ink*; and (3) a laponite/gelatin bioink without added growth factors, where TGF-β3 (10 ng/ml) was supplemented directly into the media (CDM+). The incorporation of growth factors did not affect print quality, as indicated by comparable printed filament diameters for the different groups (Fig. 4 E). Filament diameter decreased over time in all conditions (Fig. 4 E). In the CDM group, diameters were initially ∼0.45–0.50 mm and progressively declined to ∼0.30 mm by day 28 (p < 0.0001). µTs printed with the chondro-ink generated filaments with slightly smaller diameters (∼0.30–0.35 mm), with further reductions by day 28 (p < 0.001). Similarly, fibro-ink printed constructs decreased from ∼0.35–0.40 mm at baseline to ∼0.25–0.30 mm over the same period (p < 0.001). These consistent reductions likely reflect a combination of compaction imposed by the support bath and biologically driven remodeling, as growth factor release promoted µT fusion and relatively homogeneous tissue formation (Fig. 4 D). Cell viability was evaluated after 28 days of culture under all conditions. High levels of viability were observed in all printed filaments, indicating that the inclusion of laponite, sustained growth factor release, and the bioprinted microenvironment did not compromise cell survival (Fig. 4 F).

### 3.3 Co-bioprinting of growth factors and µTs supports the development of (fibro)cartilage-like tissue

The composition of the bioprinted filaments was next assessed using a range of histological biochemical assays. DNA content was comparable across groups (Fig. 5 B). By the end of the culture period, Alcian blue staining revealed sGAG deposition in all groups, though slightly reduced in the fibro-ink group (Fig. 5 A, C). Biochemical quantification confirmed that sGAG levels were significantly higher in the CDM+ and chondro-ink groups compared to the fibro-ink group (Fig. 5 E). Collagen synthesis was lower in the fibro-ink group compared to the CDM+ group (Fig. 5 D, F).

**Figure 5.**
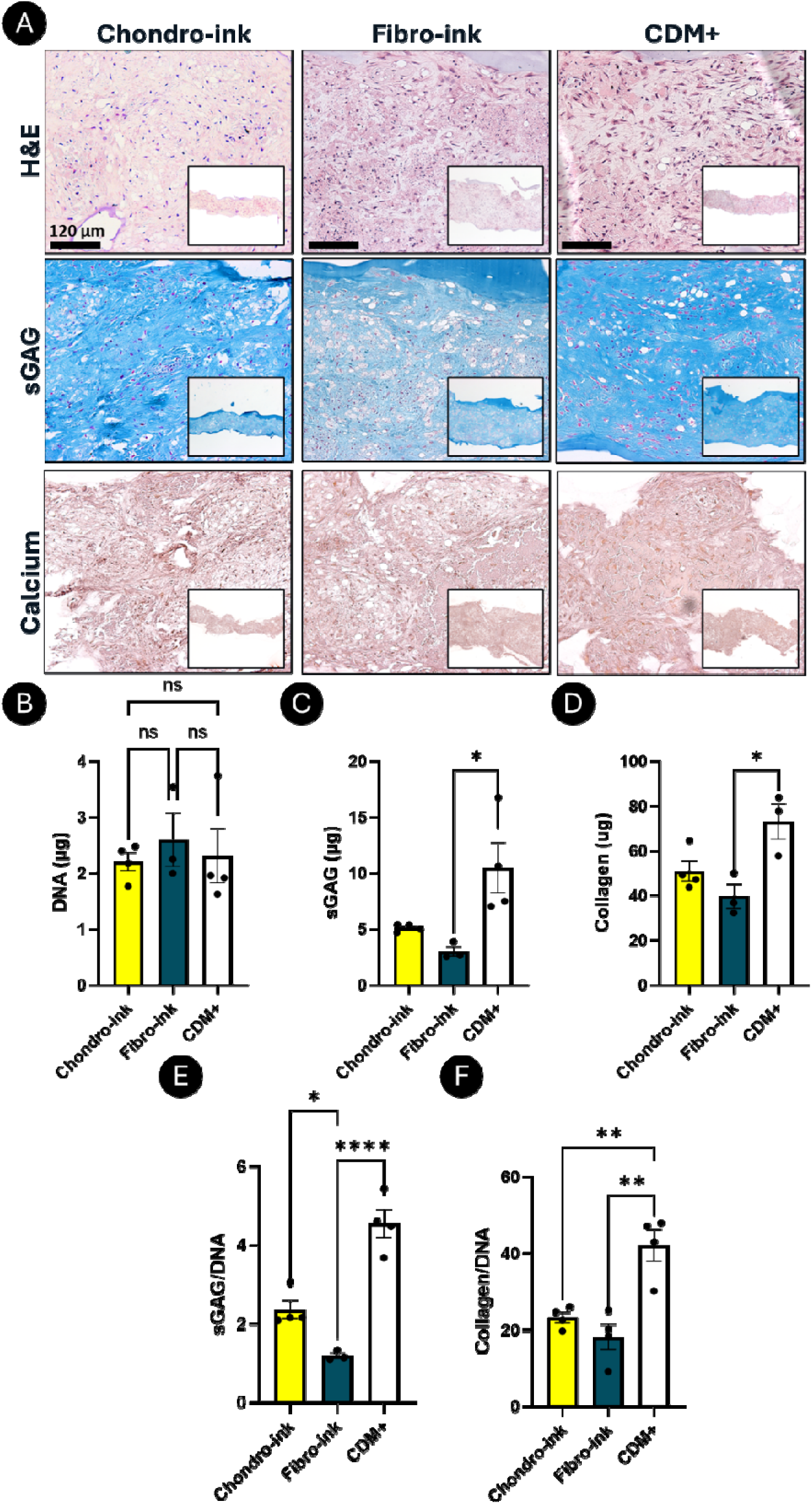
Printed µTs remodel their ECM toward a fibrocartilage phenotype within 28 days when encapsulated within a controlled release bioink. µTs were bioprinted into 1% XG-MA support bath using a gelatin-laponite bioink supplemented with the following growth factors: TGF-β3 (chondro-ink) for 4 weeks, CTGF+TGF-β3 (fibro-ink) for 4 weeks, or no growth factors in the ink but added in the medium (CDM+ group). A) Histological assessment of chondrogenic markers: H&E, Alcian Blue (sGAG), and Alizarin Red (calcium deposits). Scale bar:120 µm. B–G) Biochemical quantification: DNA (B), sGAG (C), total collagen (D), sGAG/DNA (E), collagen/DNA (F). Outliers were identified using a ROUT test. Data were analyzed by one-way ANOVA followed by Tukey’s post hoc test. Statistical significance was defined as p < 0.05 (*p < 0.05, **p < 0.01, ***p < 0.001, ns = not significant).

Collagen deposition was also assessed using Picrosirius red staining (Fig. 6 A). Notably, collagen deposition was robust in all groups, and polarized light imaging revealed collagen alignment parallel to the long axis of the printed filament in all constructs (Fig. 6 A). No major differences in collagen fiber alignment and coherency were detected between the CDM+, fibro-ink, and chondro-ink groups (Fig. 6 B, C). These findings demonstrate the potential of printing growth factor releasing inks into a XG-MA support bath to direct both the organization and composition of engineered region-specific meniscal tissues.

**Figure 6.**
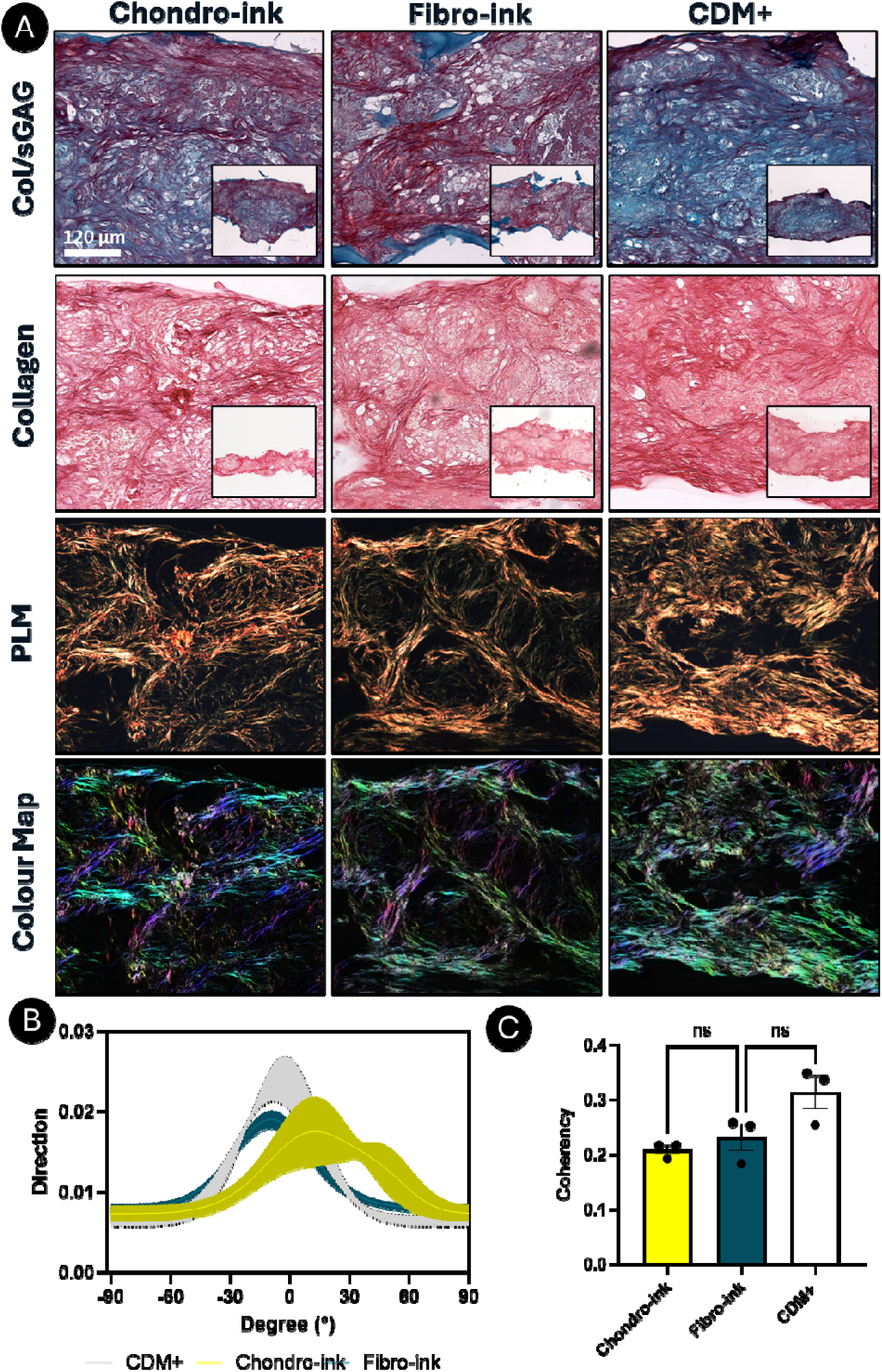
Printed µTs into laponite based bioinks exhibit distinct collagen alignment profiles under controlled release of growth factors. A) Histological staining with Picrosirius Red/Alcian Blue to visualize both collagen and sGAG deposition over 28 days, alongside polarized light microscopy (PLM) and OrientationJ-derived color maps. Color hue indicates fiber orientation: blue/cyan = 0°, pink/red = 90°. Scale bar = 120 µm. B) Average collagen fiber orientation under different growth factor regimes with laponite-mediated slow release. C) Fiber coherency, where values close to 1 represent highly aligned fibers and values near 0 represent dispersed orientations (n = 3). Scale bar = 200 µm. Statistical analysis was performed using two-way ANOVA with Tukey’s post hoc test. Statistical significance was set at p < 0.05.

### 3.4 Growth factor release and support bath confinement promote a fibrocartilage-like phenotype in printed µTs

Having shown that loading CTGF and TGF-β3 into the bioink can drive fibrocartilaginous differentiation of bioprinted µTs within the support bath, we next characterized the tissue phenotypes supported by these bioink combinations. To this end, qPCR analysis and immunohistology staining for collagen deposition were performed after 4 weeks of *in vitro* culture to assess gene expression profiles and tissue maturation associated with local growth factor delivery (Fig. 7).

**Figure 7.**
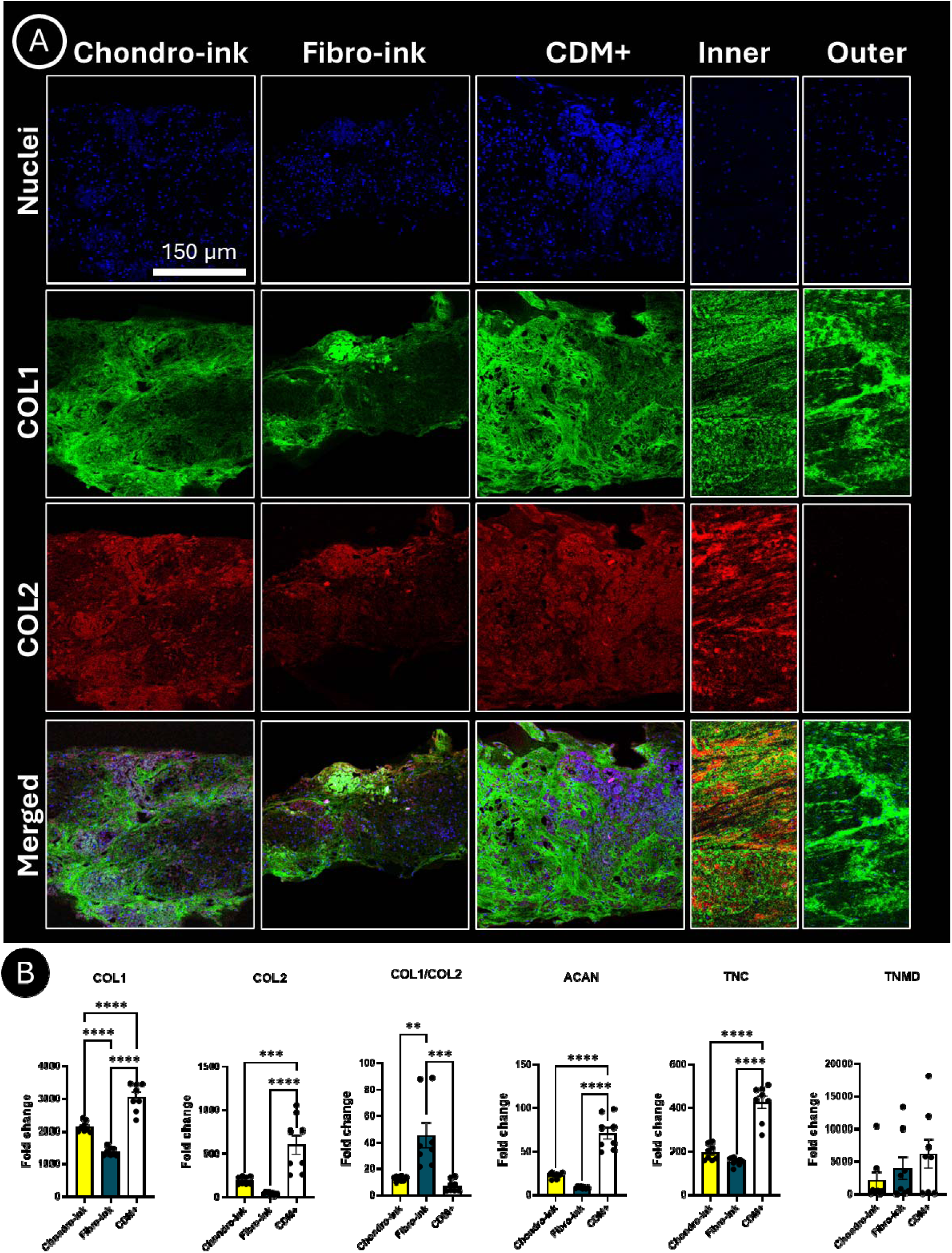
Localized growth factor delivery alters collagen deposition and gene expression in printed µTs. A) Immunohistology staining for collagen I (green) and collagen II (red) in bioprinted µTs. Native inner and outer meniscus regions are shown for comparison, highlighting regional differences in collagen deposition. Scale bar: 150 µm. B) qPCR analysis of bioprinted µTs with localized growth factor delivery, showing differential expression of chondrogenic and fibrochondrogenic markers, including collagen I (COL1), collagen II (COL2), COL1/COL2 ratio, aggrecan (ACAN), tenascin (TNC), and tenomodulin (TNMD). Data were analyzed using one-way ANOVA, with significance accepted at p < 0.05 (*p < 0.05, **p < 0.01, ***p < 0.001, ****p < 0.0001).

µTs induced within the chondro-ink stained more intensely for type II collagen, consistent with an inner meniscus phenotype (Fig. 7 A). In contrast, the µTs induced within the fibro-ink stained more intensely collagen type I but weaker collagen type II deposition, more closely resembling the outer meniscus (Fig. 7 A). Interestingly, the CDM+ and the chondro-ink groups displayed comparable staining patterns, both with high levels of collagen type I and II (Fig. 7 A). Gene expression profiles also reflected these trends. Collagen type I and II (COL1 and COL2, respectively) gene expression was highest in the CDM+ group, where TGF-β3 was added directly to the media (Fig. 7 B). Collagen type II (COL2) expression did not differ significantly between the two groups with printed growth factor delivery, but analysis of the COL1/COL2 ratio revealed a marked distinction: CDM+ and chondro-ink groups showed lower ratios, while the fibro-ink group exhibited the highest ratio, suggesting that co-stimulation with CTGF and TGF-β3 preferentially supports collagen type I over collagen type II expression (Fig. 7 B). Aggrecan (ACAN) expression was also elevated in the chondro-ink group compared to the fibro-ink group, consistent with its role in promoting a more hyaline cartilage–like phenotype (Fig. 7 B).

### 3.6 Dual-cartridge bioprinting enables precise fabrication of scaled-up regionally defined meniscal grafts

Having demonstrated that controlled release of CTGF and TGF-β3 alongside printed µTs supports fibrocartilaginous differentiation, while release of TGF-β3 alone supports a more hyaline-like phenotype, we next explored bioprinting a zonally defined meniscal graft.

Specifically, circumferential lines of µTs encapsulated in a laponite/gelatin bioink loaded with CTGF + TGF-β3 were bioprinted for the outer region of the meniscal graft (fibro-ink) (Fig. 8). For the inner region, additional circumferential lines were printed inside the outer ring, using µTs encapsulated in a laponite/gelatin bioink loaded with TGF-β3 only (chondro-ink)(Fig. 8 A). This dual-bioink approach required the use of two separate printheads, with printhead 1 depositing the fibro-ink and printhead 2 depositing the chondro-ink (Fig. 8 A-G). Microscopically, the filaments appeared well defined and closely packed, and by day 7, the µTs had already begun fusing, forming a more continuous tissue. By day 28, the gaps between filaments were further filled, indicating sustained viability and ECM deposition (Fig. 8 D). Histological analysis of the bioprinted tissues was then undertaken, where the engineered construct was divided into outer and inner regions (Fig. 8 E). The engineered inner zone stained more intensely for sGAG compared to the engineered outer zone, consistent with the effect of the chondro-ink in promoting GAG deposition, even in a scaled-up construct with the same basal media (chemically defined media without TGF-β3 and/or CTGF). Collagen staining suggested no overall differences in amount or extent of fiber alignment, in line with our previous findings. However, collagen type-specific staining showed that the engineered outer zone stained more intensely for collagen type I and less for type II, whereas the engineered inner zone stained for both collagen types I and II (Fig. 8 D).

**Figure 8.**
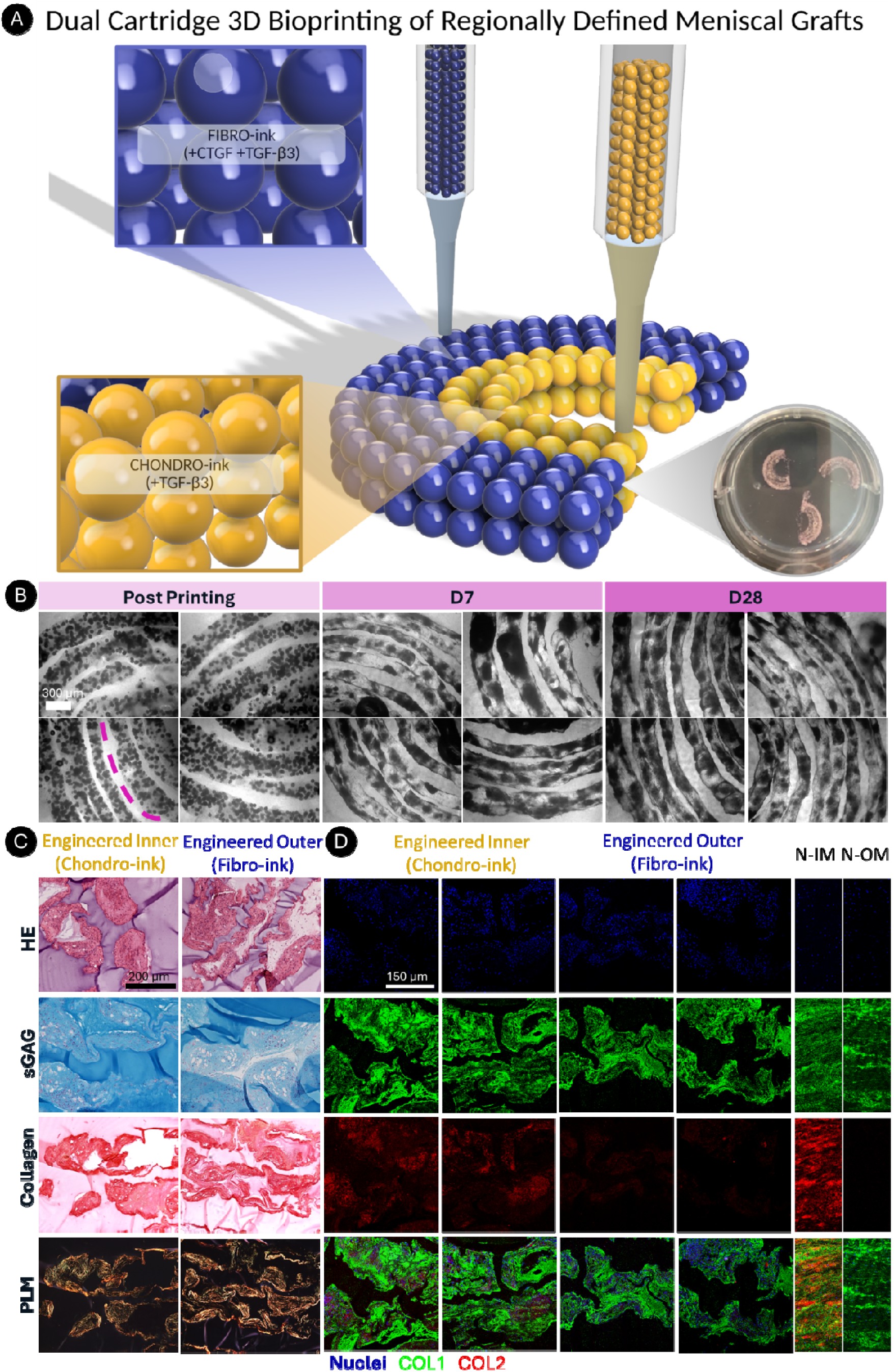
Dual-cartridge bioprinting enables precise fabrication of scaled-up grafts and region-specific constructs for meniscal regeneration. (A) Schematic of the dual-cartridge bioprinting workflow: MSC-derived µTs are deposited within an XG-MA support bath using two bioinks, a chondro-ink (+TGF-β3) and a fibro-ink (+CTGF + TGF-β3), to recapitulate meniscal zonal composition. Zonal patterning strategy: the outer zone is printed with fibro-ink (CTGF + TGF-β3) and the inner zone with chondro-ink (TGF-β3). Macroscopic image of three zonally defined bioprinted menisci immediately post-printing (day 0) in a 6-well plate. (B) Brightfield microscopy images of bioprinted filaments in the engineered inner and outer zones at day 0 and day 28; the dashed line marks the boundary between zones. Scale bar: 300 µm. (C) Histological evaluation of engineered inner and outer zones: H&E, Alcian Blue (sGAG), Picrosirius Red (collagen) and PLM. Scale bar: 200 µm. (D) Immunofluorescence for collagen I, collagen II, with DAPI for nuclei; native inner (N-IM) and outer meniscus (N-OM) shown for comparison. Scale bar: 150 µm.

Together, these results demonstrate the successful bioprinting of a biomimetic and zonally defined meniscal graft using MSC-derived µTs with spatially defined co-delivery of growth factors within a scaled-up construct.

## 4. Discussion

In this study, we successfully engineered zonally defined meniscal grafts by combining MSC-derived µTs with spatially patterned growth factor delivery through dual-cartridge bioprinting. Unlike previous scaffold-based or scaffold-free strategies that either lacked zonal fidelity or relied on meniscus-derived cells[14, 24, 37], this approach integrates a scalable, clinically accessible cell source with a dual-bioink system capable of recapitulating regional heterogeneity. To capture this heterogeneity, we expanded beyond delivery of TGF-β3 alone and incorporated CTGF in laponite/gelatin bioinks to spatially pattern growth factors in bioprinted constructs. It was found that bioinks enriched with TGF-β3 alone modulated matrix deposition of printed µTs toward a phenotype resembling the inner region of the native meniscus. Conversely, bioinks containing both TGF-β3 and CTGF promoted matrix deposition more closely associated with the outer region of the native meniscus. As a step toward engineering a scaled-up meniscal graft that recapitulate both inner and outer regions, we successfully bioprinted a zone-specific construct displaying distinct collagen deposition patterns, with collagen type II enriched in the inner zone and collagen type I enriched in the outer zone of the bioprinted graft.

Previous studies had shown that CTGF enhances collagen type I deposition while suppressing collagen type II, whereas TGF-β3 stimulation promotes both sGAG and collagen type II accumulation[16, 18, 38]. Based on this rationale, we first evaluated different growth factor stimulation regimes in self-assembled µT constructs to determine their suitability for zonal meniscus bioprinting. MSC derived µTs were allowed to fuse in agarose molds under five conditions: A)TGF-β3 alone, B) CTGF alone, C) sequential CTGF→TGF-β3, D) sequential TGF-β3→CTGF, and E) CTGF + TGF-β3. This 3D µT system allowed us to directly assess phenotypical responses to different growth factor stimulation regimes, which can differ from those reported in 2D cultures or cell-laden hydrogels [18]. It was found that CTGF stimulation alone was insufficient for promoting fibrochondrogenic differentiation, as evident by relatively low levels of sGAG and collagen deposition. However, when combined with TGF-β3, CTGF modulated matrix composition to more closely mimic that observed in the outer zone of the meniscus, which contains relatively low levels of sGAGs but which is rich in type I collagen [39]. Immunostaining further confirmed that TGF-β3 promoted strong collagen type I and II expression, while CTGF shifted the balance toward higher collagen I and markedly reduced collagen II deposition. Overall, the growth factors regimes produced responses that were broadly consistent with previous studies, with one exception. In a study by Lee et al. (2014), the sequential CTGF to TGF-β3 supported a more fibrochondrogenic phenotype, with positive staining for sGAG, collagen type I and II[18]. In contrast, in the present study, collagen type II staining was most prominent in the TGF-β3 only group compared with all other conditions. This discrepancy may be explained by differences in the 3D culture systems used. In our study, MSCs were cultured as a scaffold-free self-assembled µT system, whereas Lee et al. used MSCs embedded within a fibrin hydrogel. These differences in culture architecture and matrix environment may influence growth factor responsiveness and subsequent matrix deposition. Overall, these observations provided a clear rationale for selecting TGF-β3 alone to support an inner zone phenotype, and CTGF + TGF-β3 for the outer zone in subsequent bioprinting studies.

The bioprinting of regionally defined meniscal grafts required the development of bioinks to control the release of these growth factors to fusing µTs. It was found that the incorporation of 0.5% laponite into gelatin-based inks supported sustained release of both CTGF and TGF-β3 over 4 weeks of culture. This is in agreement with previous studies that found laponite supported the controlled release of BMP-2 over 14 days when incorporated into alginate/methylcellulose inks [40]. While local growth factor release from our fibro-ink and chondro-ink supported zone-specific meniscal phenotypes, it was noted that total matrix synthesis in this group still lagged that observed in control constructs where the growth factor was supplemented directly into the culture media. This suggests that further optimization of these bioinks may be possible, for example by modulating the total concentration of growth factor that is loaded into the inks prior to printing. Furthermore, from a translational perspective, the use of laponite as a component of the bioink needs careful consideration. While laponite has been widely explored in TE due to its favorable rheological properties and ability to bind and release growth factors [40, 41], it is not currently approved as an implantable material for clinical use. Therefore, prior to clinical translation, its safety, degradation profile, and long-term biological interactions would need to be thoroughly evaluated in vivo and assessed through appropriate regulatory pathways. Strategies such as reducing laponite content, ensuring its clearance over time, or replacing it with clinically approved alternatives may be required. However, its use in this study provides a valuable proof-of-concept for controlled growth factor delivery within bioprinted µT-based constructs, which can inform the development of more clinically translatable systems.

The engineering of functional meniscal grafts requires not only recapitulating the zonal composition of the native tissue, but also key aspects of the structure of the extracellular matrix that is integral to its function. We previously found that the mechanical properties of support baths regulate both tissue phenotype and the (re)modelling of bioprinted µTs, with intermediate stiffness (1% XG-MA) support baths supporting the development of structurally aligned fibrocartilage-like tissue [25]. Here we extend these findings and demonstrate that such physically confining support baths also support the development of a preferentially aligned collagen matrix when growth factors are loaded directly into a supporting ink. In doing so it was possible to local regulate both tissue phenotype (via local growth factor delivery), as well as matrix organization (via the physical constraints provided by the support bath). Collectively these findings point to the clear importance of providing appropriate physical boundaries to developing tissues in order to guide their development and organization.

By leveraging the self-assembly potential of µTs [21, 42] and the controlled growth factor release properties of the laponite-based bioinks [40], we were able to bioprint zonally defined meniscal constructs that reproduced key features of the native tissue. All printed groups showed high cell viability. Constructs cultured with the fibro-ink exhibited fibrocartilaginous characteristics consistent with the outer meniscus, whereas the chondro-ink group induced a more hyaline-like phenotype consistent with the inner zone. A major novelty of this work lies in the successful use of dual-cartridge bioprinting to achieve 1) homogeneous µT deposition and relatively high resolution with the use of 2 different printheads and 2) zonal µT and growth factor patterning within a single meniscal graft. Previous studies employing growth factor–loaded microparticles [43–45], homogeneous bioinks [46, 47], or meniscus progenitor cells [37] [48–51] achieved only partial zonal organization and/or were constrained to smaller scales. Here, the integration of two distinct bioinks enabled precise spatial patterning of µTs and growth factor combinations supportive of either inner or outer meniscus phenotypes, while maintaining filament fidelity and tissue compactness at the macroscale. Histological analysis of scaled-up constructs confirmed effective spatial delivery of growth factors: the engineered inner zone displayed higher sGAG accumulation together with strong collagen type II and collagen type I deposition, while the engineered outer zone exhibited lower sGAG levels, negligible collagen type II and robust collagen type I deposition.

These results demonstrate that laponite-based growth factor delivery can successfully promote region-specific differentiation in large, engineered constructs—a persistent challenge in TE. Compared with our previous studies[25, 52], this study marks a significant step forward by translating small-scale findings into large, architecturally defined grafts that preserve biomimicry during in vitro culture. Although collagen deposition improved with combined growth factor regimes, total collagen content remained below native levels, and collagen fiber anisotropy has not yet been fully recapitulated. Future work could explore mechanical stimulation and extended culture periods to enhance collagen maturation and alignment, as well as evaluate the mechanical properties of the grafts under physiologically relevant loading. Ultimately, in vivo validation will be required to assess tissue integration and long-term functionality.

In summary, this work introduces a novel strategy to advance the field for the biofabrication of zonally defined meniscal grafts by combining MSC-derived µTs with dual-bioink delivery of growth factors. By uniting scalable cell sources, precise spatial patterning in physically constraining support baths and sustained but localized growth factor presentation, this approach advances beyond prior scaffold-based or single-bioink methods and establishes a robust platform for developing clinically relevant meniscal replacements.

## 5. Conclusions

This study establishes a framework for engineering zonally defined meniscal grafts by integrating MSC-derived µTs with dual-cartridge bioprinting and laponite-based growth factor delivery. The constructs successfully recreated key biochemical gradients of the native tissue: the inner zone displayed higher sGAG and collagen type II deposition, while the outer zone exhibited lower sGAG content and predominantly collagen type I, consistent with its fibrocartilaginous character.

Importantly, these zonal differences were preserved in scaled-up grafts despite shared media conditions, underscoring the effectiveness of localized growth factor presentation. While total collagen deposition and fiber anisotropy remain below native benchmarks, this approach advances beyond previous scaffold-based or single-bioink strategies by combining scalable cell sources, localized biochemical and biophysical guidance, and spatial fidelity in a single graft. Taken together, these findings represent a significant step toward the development of clinically relevant, biomimetic meniscal implants.

## 7. Acknowledgements

We would like to thank Dr Megan Canavan from Trinity College Dublin for the SEM imaging. Schematic diagrams of the graphical abstract, Figures 1 and 2 were created with BioRender.com and Autodesk Fusion 360 software by F.D.S. This work was funded by the European Research Council (ERC, 4D-BOUNDARIES #101019344)

## References

[1] L.S. Lohmander, P.M. Englund, L.L. Dahl, E.M. Roos, The long-term consequence of anterior cruciate ligament and meniscus injuries: osteoarthritis, Am J Sports Med 35(10) (2007) 1756–69.

[2] L.D. Bennett, J.C. Buckland-Wright, Meniscal and articular cartilage changes in knee osteoarthritis: a cross-sectional double-contrast macroradiographic study, Rheumatology (Oxford) 41(8) (2002) 917–23.

[3] J.T. Badlani, C. Borrero, S. Golla, C.D. Harner, J.J. Irrgang, The effects of meniscus injury on the development of knee osteoarthritis: data from the osteoarthritis initiative, Am J Sports Med 41(6) (2013) 1238–44.

[4] X. Liu, H. Huang, W. Yin, S. Ren, Q. Rong, Y. Ao, Anterior cruciate ligament deficiency combined with lateral and/or medial meniscal injury results in abnormal kinematics and kinetics during level walking, Proc Inst Mech Eng H 234(1) (2020) 91–99.

[5] P.H. Lento, V. Akuthota, Meniscal injuries: A critical review, J Back Musculoskelet Rehabil 15(2) (2000) 55–62.

[6] B. Bilgen, C.T. Jayasuriya, B.D. Owens, Current Concepts in Meniscus Tissue Engineering and Repair, Adv Healthc Mater 7(11) (2018) e1701407.

[7] B. Vundelinckx, J. Vanlauwe, J. Bellemans, Long-term Subjective, Clinical, and Radiographic Outcome Evaluation of Meniscal Allograft Transplantation in the Knee, Am J Sports Med 42(7) (2014) 1592–9.

[8] N.A. Smith, N. MacKay, M. Costa, T. Spalding, Meniscal allograft transplantation in a symptomatic meniscal deficient knee: a systematic review, Knee Surg Sports Traumatol Arthrosc 23(1) (2015) 270–9.

[9] E. Rath, J.C. Richmond, The menisci: basic science and advances in treatment, Br J Sports Med 34(4) (2000) 252–7.

[10] P. Ghosh, T.K. Taylor, The knee joint meniscus. A fibrocartilage of some distinction, Clin Orthop Relat Res (224) (1987) 52–63.

[11] H.E. Kambic, C.A. McDevitt, Spatial organization of types I and II collagen in the canine meniscus, J Orthop Res 23(1) (2005) 142–9.

[12] P.G. Bullough, L. Munuera, J. Murphy, A.M. Weinstein, The strength of the menisci of the knee as it relates to their fine structure, J Bone Joint Surg Br 52(3) (1970) 564–7.

[13] K. Nakata, K. Shino, M. Hamada, T. Mae, T. Miyama, H. Shinjo, S. Horibe, K. Tada, T. Ochi, H. Yoshikawa, Human meniscus cell: characterization of the primary culture and use for tissue engineering, Clin Orthop Relat Res (391 Suppl) (2001) S208–18.

[14] B.M. Baker, A.S. Nathan, G.R. Huffman, R.L. Mauck, Tissue engineering with meniscus cells derived from surgical debris, Osteoarthritis Cartilage 17(3) (2009) 336–45.

[15] J. Baek, X. Chen, S. Sovani, S. Jin, S.P. Grogan, D.D. D’Lima, Meniscus tissue engineering using a novel combination of electrospun scaffolds and human meniscus cells embedded within an extracellular matrix hydrogel, J Orthop Res 33(4) (2015) 572–83.

[16] S. Tarafder, J. Gulko, D. Kim, K.H. Sim, S. Gutman, J. Yang, J.L. Cook, C.H. Lee, Effect of dose and release rate of CTGF and TGFβ3 on avascular meniscus healing, J Orthop Res 37(7) (2019) 1555–1562.

[17] C.H. Lee, B. Shah, E.K. Moioli, J.J. Mao, CTGF directs fibroblast differentiation from human mesenchymal stem/stromal cells and defines connective tissue healing in a rodent injury model, J Clin Invest 120(9) (2010) 3340–9.

[18] C.H. Lee, S.A. Rodeo, L.A. Fortier, C. Lu, C. Erisken, J.J. Mao, Protein-releasing polymeric scaffolds induce fibrochondrocytic differentiation of endogenous cells for knee meniscus regeneration in sheep, Sci Transl Med 6(266) (2014) 266ra171.

[19] L.C. Ionescu, G.C. Lee, K.L. Huang, R.L. Mauck, Growth factor supplementation improves native and engineered meniscus repair in vitro, Acta Biomater 8(10) (2012) 3687–94.

[20] R. Burdis, D.J. Kelly, Biofabrication and bioprinting using cellular aggregates, microtissues and organoids for the engineering of musculoskeletal tissues, Acta Biomater 126 (2021) 1–14.

[21] V. Mironov, R.P. Visconti, V. Kasyanov, G. Forgacs, C.J. Drake, R.R. Markwald, Organ printing: tissue spheroids as building blocks, Biomaterials 30(12) (2009) 2164–74.

[22] G.S. Kronemberger, K. Chattahy, F.D. Spagnuolo, A.S. Karam, D.J. Kelly, Bioassembly of Region-Specific Fibrocartilage Microtissues to Engineer Zonally Defined Meniscal Grafts, Adv Healthc Mater (2025) e02208.

[23] W. Niu, W. Guo, S. Han, Y. Zhu, S. Liu, Q. Guo, Cell-Based Strategies for Meniscus Tissue Engineering, Stem Cells Int 2016 (2016) 4717184.

[24] P. Angele, B. Johnstone, R. Kujat, J. Zellner, M. Nerlich, V. Goldberg, J. Yoo, Stem cell based tissue engineering for meniscus repair, J Biomed Mater Res A 85(2) (2008) 445–55.

[25] F.D. Spagnuolo, G.S. Kronemberger, D.J. Kelly, Bioprinting of Microtissues Within Mechanically Tunable Support Baths to Engineer Anisotropic Musculoskeletal Tissues, Adv Sci (Weinh) 13(18) (2026) e09313.

[26] P. Bordes, E. Pollet, L. Averous, Nano-biocomposites: Biodegradable polyester/nanoclay systems, Progress in Polymer Science 34(2) (2009) 125–155.

[27] A.K. Gaharwar, L.M. Cross, C.W. Peak, K. Gold, J.K. Carrow, A. Brokesh, K.A. Singh, 2D Nanoclay for Biomedical Applications: Regenerative Medicine, Therapeutic Delivery, and Additive Manufacturing, Adv Mater 31(23) (2019) e1900332.

[28] L.M. Cross, J.K. Carrow, X. Ding, K.A. Singh, A.K. Gaharwar, Sustained and Prolonged Delivery of Protein Therapeutics from Two-Dimensional Nanosilicates, ACS Appl Mater Interfaces 11(7) (2019) 6741–6750.

[29] F.D. Spagnuolo, G.S. Kronemberger, K.J. Storey, D.J. Kelly, The maturation state and density of human cartilage microtissues influence their fusion and development into scaled-up grafts, Acta Biomater 194 (2025) 109–121.

[30] A.M. Mackay, S.C. Beck, J.M. Murphy, F.P. Barry, C.O. Chichester, M.F. Pittenger, Chondrogenic differentiation of cultured human mesenchymal stem cells from marrow, Tissue Eng 4(4) (1998) 415–28.

[31] A.H. Huang, A. Stein, R.S. Tuan, R.L. Mauck, Transient exposure to transforming growth factor beta 3 improves the mechanical properties of mesenchymal stem cell-laden cartilage constructs in a density-dependent manner, Tissue Eng Part A 15(11) (2009) 3461–72.

[32] J. Herwig, E. Egner, E. Buddecke, Chemical changes of human knee joint menisci in various stages of degeneration, Ann Rheum Dis 43(4) (1984) 635–40.

[33] S.G. Patrício, L.R. Sousa, T.R. Correia, V.M. Gaspar, L.S. Pires, J.L. Luís, J.M. Oliveira, J.F. Mano, Freeform 3D printing using a continuous viscoelastic supporting matrix, Biofabrication 12(3) (2020) 035017.

[34] M. Ahn, W.W. Cho, H. Lee, W. Park, S.H. Lee, J.W. Back, Q. Gao, G. Gao, D.W. Cho, B.S. Kim, Engineering of Uniform Epidermal Layers via Sacrificial Gelatin Bioink-Assisted 3D Extrusion Bioprinting of Skin, Adv Healthc Mater 12(27) (2023) e2301015.

[35] Y. Zhang, M. Chen, Z. Dai, H. Cao, J. Li, W. Zhang, Sustained protein therapeutics enabled by self-healing nanocomposite hydrogels for non-invasive bone regeneration, Biomater Sci 8(2) (2020) 682–693.

[36] D. Kilian, S. Cometta, A. Bernhardt, R. Taymour, J. Golde, T. Ahlfeld, J. Emmermacher, M. Gelinsky, A. Lode, Core-shell bioprinting as a strategy to apply differentiation factors in a spatially defined manner inside osteochondral tissue substitutes, Biofabrication 14(1) (2022).

[37] X. Barceló, K.F. Eichholz, I.F. Gonçalves, O. Garcia, D.J. Kelly, Bioprinting of structurally organized meniscal tissue within anisotropic melt electrowritten scaffolds, Acta Biomater 158 (2023) 216–227.

[38] W. He, Y.J. Liu, Z.G. Wang, Z.K. Guo, M.X. Wang, N. Wang, Enhancement of meniscal repair in the avascular zone using connective tissue growth factor in a rabbit model, Chin Med J (Engl) 124(23) (2011) 3968–75.

[39] U.P. Palukuru, C.M. McGoverin, N. Pleshko, Assessment of hyaline cartilage matrix composition using near infrared spectroscopy, Matrix Biol 38 (2014) 3–11.

[40] F.E. Freeman, P. Pitacco, L.H.A. van Dommelen, J. Nulty, D.C. Browe, J.Y. Shin, E. Alsberg, D.J. Kelly, 3D bioprinting spatiotemporally defined patterns of growth factors to tightly control tissue regeneration, Sci Adv 6(33) (2020) eabb5093.

[41] F.E. Freeman, D.J. Kelly, Tuning Alginate Bioink Stiffness and Composition for Controlled Growth Factor Delivery and to Spatially Direct MSC Fate within Bioprinted Tissues, Sci Rep 7(1) (2017) 17042.

[42] J.Y. Schell, B.T. Wilks, M. Patel, C. Franck, V. Chalivendra, X. Cao, V.B. Shenoy, J.R. Morgan, Harnessing cellular-derived forces in self-assembled microtissues to control the synthesis and alignment of ECM, Biomaterials 77 (2016) 120–9.

[43] O. Jeon, H. Park, J.K. Leach, E. Alsberg, Biofabrication of engineered tissues by 3D bioprinting of tissue specific high cell-density bioinks, bioRxiv (2024).

[44] E. Zhou, P. He, Z. Yang, C. Li, G. Fang, J. Wu, W. Zhuang, H. Sang, 3D-printed GelMA-Alginate microsphere scaffold with staged dual-growth factor release for enhanced bone regeneration, Mater Today Bio 35 (2025) 102422.

[45] W. Fang, M. Yang, M. Liu, Y. Jin, Y. Wang, R. Yang, Y. Wang, K. Zhang, Q. Fu, Review on Additives in Hydrogels for 3D Bioprinting of Regenerative Medicine: From Mechanism to Methodology, Pharmaceutics 15(6) (2023).

[46] A. Bandyopadhyay, B. Ghibhela, S. Shome, S. Hoque, S.K. Nandi, B.B. Mandal, Photo-Polymerizable Autologous Growth-Factor Loaded Silk-Based Biomaterial-Inks toward 3D Printing-Based Regeneration of Meniscus Tears, Adv Biol (Weinh) 8(5) (2024) e2300710.

[47] E. Stocco, A. Porzionato, E. De Rose, S. Barbon, R. De Caro, V. Macchi, Meniscus regeneration by 3D printing technologies: Current advances and future perspectives, J Tissue Eng 13 (2022) 20417314211065860.

[48] J. Korpershoek, M. Rikkers, T.S. De Windt, M.A. Tryfonidou, D.B. Saris, L.A. Vonk, Progenitor cells with high chondrogenic potential are present in the adult human meniscus, Osteoarthritis and Cartilage 28 (2020) S205–S206.

[49] J.V. Korpershoek, M. Rikkers, T.S. de Windt, M.A. Tryfonidou, D.B.F. Saris, L.A. Vonk, Selection of Highly Proliferative and Multipotent Meniscus Progenitors through Differential Adhesion to Fibronectin: A Novel Approach in Meniscus Tissue Engineering, Int J Mol Sci 22(16) (2021).

[50] G. Pattappa, F. Reischl, J. Jahns, R. Schewior, S. Lang, J. Zellner, B. Johnstone, D. Docheva, P. Angele, Fibronectin Adherent Cell Populations Derived From Avascular and Vascular Regions of the Meniscus Have Enhanced Clonogenicity and Differentiation Potential Under Physioxia, Front Bioeng Biotechnol 9 (2021) 789621.

[51] X. Barceló, K. Eichholz, I. Gonçalves, G.S. Kronemberger, A. Dufour, O. Garcia, D.J. Kelly, Bioprinting of scaled-up meniscal grafts by spatially patterning phenotypically distinct meniscus progenitor cells within melt electrowritten scaffolds, Biofabrication 16(1) (2023).

[52] G.S. Kronemberger, F.D. Spagnuolo, A.S. Karam, K. Chattahy, K.J. Storey, D.J. Kelly, Rapidly Degrading Hydrogels to Support Biofabrication and 3D Bioprinting Using Cartilage Microtissues, ACS Biomater Sci Eng 10(10) (2024) 6441–6450.

